# Chemerin Induced by *Treponema pallidum* Predicted Membrane Protein Tp0965 Mediates Endothelial Dysfunction via Activating MAPK Signaling Pathway

**DOI:** 10.1101/423160

**Authors:** Rui-Li Zhang, Li-Jia Yang, Qian-Qiu Wang

## Abstract

Chemerin, a chemoattractant protein, is involved in endothelial dysfunction and vascular inflammation in pathological conditions. In a recent study, we observed the upregulation of chemerin in endothelial cells following *in vitro* treatment with *T. pallidum*. Here, we investigated the role of chemerin in endothelial cells dysfunction induced by the *T. pallidum* predicted membrane protein Tp0965. Following stimulation of human umbilical vein endothelial cells (HUVECs) with Tp0965, chemerin and its ChemR23 receptor were up-regulated, companied with elevated expression of TLR2. Furthermore, chemerin from HUVECs activated endothelial cells via chemerin/ChemR23 signaling in an autocrine/paracrine manner, characterized by upregulated expression of ICAM-1, E-selectin and MMP-2. Activation of endothelial cells depended on the MAPK signaling pathway. In addition, Tp0965-induced chemerin promoted monocytes migration to endothelial cells, also via chemerin/ChemR23 pathway. The RhoA/ROCK signaling pathway was also involved in monocytes migration in response to chemerin/ChemR23. Our results highlight the role of Tp0965-induced chemerin in endothelial cells dysfunction, which contributes to the immunopathogenesis of vascular inflammation of syphilis.

**Author summary:** *Treponema pallidum* is the spirochete of syphilis, which causes a chronic system inflammation. Endothelium damage caused by this bacterium is the key step in the systemic dissemination and pathophysiology of syphilis, particularly cardiovascular syphilis and neurosyphilis. In this study, we show a novel molecular mechanism of endothelium damage induce by *Treponema pallidum* predicted membrane protein Tp0965. Chemerin is a recently identified adipocytokine and chemoattractant protein with a crucial role in endothelial dysfunction and vascular inflammation in pathological conditions. Our data show that Tp0965 up-regulated the expression of chemerin and its ChemR23 receptor by endothelial cells in vitro. Furthermore, chemerin from HUVECs activated endothelial cells via chemerin/ChemR23 signaling in an autocrine/paracrine manner and depended on the MAPK signaling pathway. In addition, Tp0965-induced chemerin promoted monocytes migration to endothelial cells, also via chemerin/ChemR23 pathway. The RhoA/ROCK signaling pathway was also involved in monocytes migration in response to chemerin/ChemR23. These findings contribute to the immunopathogenesis of vascular inflammation of syphilis.

## Introduction

*Treponema pallidum* subsp. *pallidum* (*T. pallidum*) is the spirochete of syphilis, a sexually transmitted infection that remains an important public health problem around the world [1]. Infection by this bacterium causes a chronic system inflammation, and if untreated can seriously and irreversibly damage the nervous and cardiovascular system. Cardiovascular syphilis is associated with multiple clinical syndromes, of which aortic aneurysms, aortic insufficiency, and coronary artery stenosis are the most common [2-3]. These clinical presentations are similar to other, more common varieties of cardiac disease. At the beginning of the inflammatory process following infection by *T. pallidum*, the insult is primarily to the small nutrient vessels of the aorta, but by later stage all three layers of aortic wall are affected. The overlying aortic intima becomes diffusely diseased, with atherosclerotic changes occurring across virtually the entire intimal surface of the affected aorta. The calcification accompanying these complex atherosclerotic plaques accounts for the eggshell calcification. Characteristic histologic changes consist of infiltration by lymphocytes and plasma cells, suggesting that the condition has an immunologic basis. However, because we lack an ideal animal model of syphilis to mimic the invading progress of *T. pallidum*, the pathogenesis of these inflammatory processes are largely uncharacterized to date.

Based on an expanded corpus of molecular immunological observations, research focused on bacteria structure and physiology of bacteria made considerable progress over the past two decades [4-6]. However, investigators found that *T. pallidum* outer membrane, unlike other Gram-negative bacteria, is fragile and contains limited antigens. There is now a consensus that vessel inflammation as well as other tissue damage result from the host immune response elicited by antigen by inner membrane and periplasm of *T. pallidum*. To date, 116 proteins of *T. pallidum* have been predicted as membrane localized proteins [7]. However, the immunological characteristics of most of these membrane proteins remain unknown, and only a few membrane proteins were reported in previous studies [8-9]. Proteome array analysis predicted that Tp0965 located in inner membrane of *T. pallidum.* Tp0965 is a putative membrane fusion protein and belongs to efflux transporter RND family MFP subunit [7]. Our and others previous study suggested that recombinant protein Tp0965 showed excellent antigenicity and immunoreactivity [10-11].

Endothelial dysfunction plays key roles in vascular intimal inflammation by initiating the formation of atherosclerotic plaques and thrombosis [12]. It is also the key step of systemic dissemination and pathophysiological basic of syphilis [13-14]. We investigated the effect of Tp0965 as well as Tp17 on endothelial cells. We found that the two recombinant proteins could induce endothelial dysfunction characterized by upregulated expression of proinflammatory cytokines, as well as enhanced transendothelial migration and adhesion of monocytes to endothelial cells [15-16]. These data suggest that Tp17 and Tp0965 are involved in vessel inflammation in *T. pallidum* persistent invasion. In addition, we found that chemerin, a recently identified adipocytokine and chemoattractant protein with a crucial role in inflammation and adipocytes metabolism [17-18], is highly expressed in endothelial cells after *in vitro* treatment with *T. pallidum*. As a chemoattractant, chemerin can induce recruitment of leukocytes to inflammation sites and binding to adhesion molecules, thereby contributing to leukocyte activation [19]. Accordingly, the role of chemerin in vascular dysfunction has attracted increasing interest. Several teams reported the involvement of chemerin in multiple inflammatory disorders, such as cardiovascular disease, multiple sclerosis, lupus, psoriasis, and especially atherosclerosis [20-23].

Given the effect of chemerin and predicted membrane proteins of *T. pallidum* on vascular endothelial cells, we stimulated human umbilical vein endothelial cells (HUVECs) with recombinant Tp0965 and found that this treatment increased chemerin expression. With the aforementioned, we hypothesized that Tp0965 contributes to endothelial dysfunction by inducing chemerin expression. In order to test this hypothesis, we investigated the role of chemerin in endothelial cells dysfunction and migration of monocytes to endothelial cells, as well as the pathways involved. The data revealed that Tp0965-induced chemerin activated endothelial cells in a MAPK signaling dependent manner and promoted monocytes migration through chemerin/ChemR23 signaling pathway.

## Materials and Methods

### Ethics Statement

The research was ethically conducted under the protocol number 2016-050 approved by the Institutional Ethics Committee of Nanjing Medical University (Nanjing, China).

### Cell Culture

Both of HUVECs and THP-1 cells were obtained from the lab of Dr. Qian-Qiu Wang (Chinese Academy of Medical Sciences). HUVECs were cultured in complete EBM-2 culture media (Gibco, Grand Island, NY, containing 0.2% bovine brain extract, 5 ng/ml human EGF, 10 mM L-glutamine,1 mg/mL hydrocortisone, 2% FBS, and 0.5% streptomycin.) and used for experiments between passage 3 and 6. THP-1 cells were cultured in RPMI 1640 medium (Gibco, Grand Island, NY) containing 10% FBS, 0.5% streptomycin, and 0.05 mM 2-mercaptoethanol. All cells were cultured at 37°C in a humidified, 5% CO_2_ atmosphere.

### Construction of plasmids and cell transfection

Plasmids expressing shRNA against TLR2(shTLR2), TLR4 (shTLR4), chemerin (shchemerin) and ChemR23 (shChemR23) were designed and constructed by Zoonbio Co (Nanjing, Jiangsu, China). Transfections of HUVEC and THP-1 cells were performed with Lipofectamine 2000 (Invitrogen, Carlsbad, CA, USA). Briefly, cells were seeded into 6-well plates and incubated with vector supernatants for 6h, and then the culture medium was removed and replaced with fresh EBM-2. After an additional 48h culture, the cells were harvested and lysed for extraction of total RNA and protein. To confirm that the transduction was successful, mRNA and protein expression levels were determined by quantitative reverse transcriptase–polymerase chain reaction (qRT-PCR) and western blot. Sequences of the primers used in this study are listed in Table 1. All of the primers were designed and synthesized by Zoonbio Co.

**Table 1.**
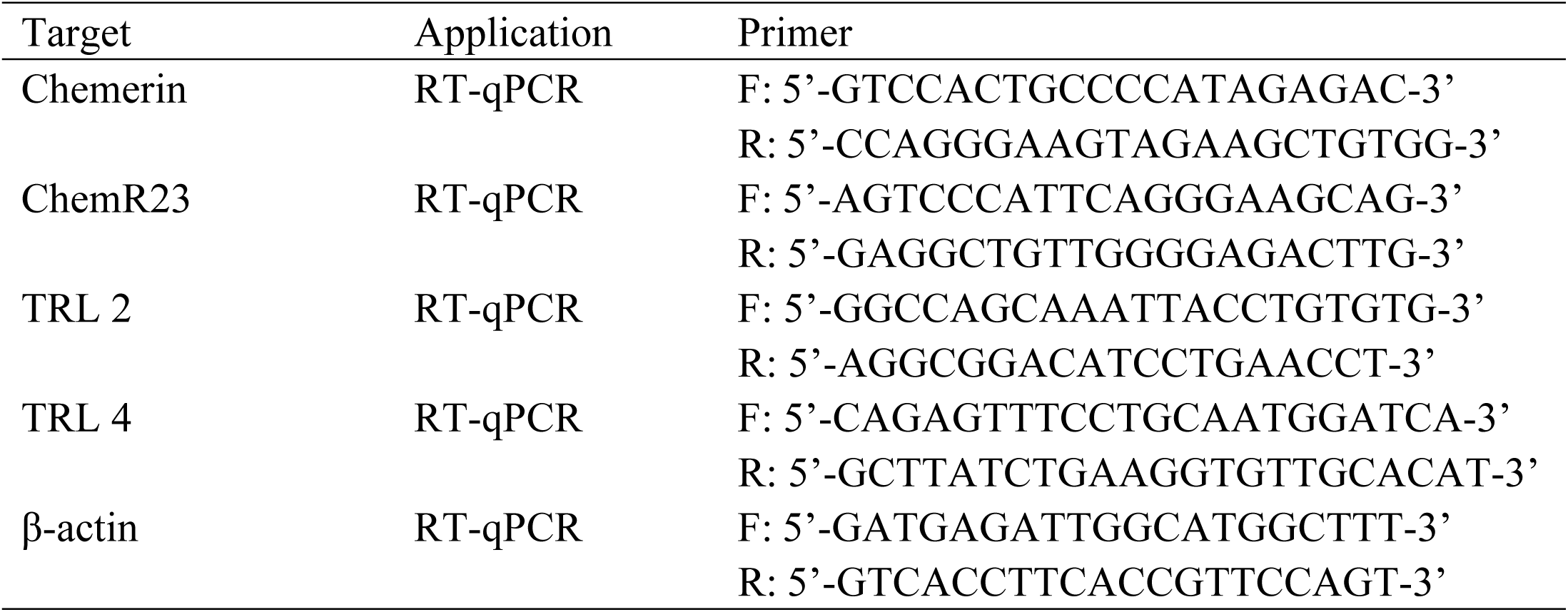
The sequences of specific primers for qPCR.

### Western blot

Western blot analysis was performed as previously described [13]. Briefly, the total protein extracted from lysed cells was electrophoresed on 10% SDS-polyacrylamide gels under denaturing conditions, followed by electrotransfer to polyvinylidene fluoride (PVDF) membranes. After blocking with 5% non-fat milk diluted in Tris buffered saline tween (TBST), the membrane was incubated with the primary antibodies purchased from Sigma-Aldrich (St Louis, MO, USA). Thereafter, the membrane was incubated with the secondary antibody labeled with horseradish peroxidase (HRP) (Sigma-Aldrich, St Louis, MO, USA). After the member was washed three times with Tris buffered saline, immunoreactive binding was detected with prepared chemiluminescence (ECL). Densitometry analysis was performed with the image analysis software (Molecular Imager Gel Doc^™^, Bio-RAD, Hercules, CA, USA) and β-actin served as a control.

### ELISA

HUVECs were seeded in 96-well plates and cultured in EBM-2 media. After treatment with Tp0965 or recombinant chemerin, culture supernatants were collected. The secreted levels of MMP-2, ICAM-1, and E-selectin were determined using enzyme-linked immunosorbent assay (ELISA) kits (R&D Systems, Minneapolis, MN, USA). Finally, the values were calculated on the basis of standard curve constructed for each assay.

### qRT-PCR

Total RNA was isolated from HUVECs using the TRIzol reagent (Invitrogen, Carlsbad, CA, USA) and cDNA was synthesized from 1 µg total RNA using the RETROscript reverse transcription kit (Invitrogen, Carlsbad, CA, USA) according to the manufacturer’s instructions. Real-time RT-PCR was performed with FastStart SYBR Green Master mix and an ABI StepOne Plus Sequence Detection System (Applied Biosystems) as previously described [13]. mRNA levels of targeted genes were determined with the ΔΔCt method and normalized against the corresponding levels of β-actin mRNA.

### Immunofluorescence assay (IFA)

Cells were plated on cover slips in 24-well plates and incubated at 37°C overnight for attachment. After three washes in PBS, cells were fixed in 5% paraformaldehyde for 1h at room temperature, and then blocked for 1h at room temperature with blocking buffer (PBS containing 0.5% BSA and 5% goat serum). Next, cells were incubated using the primary antibody (1:100 dilution) at 4°C overnight, followed by incubation with the secondary antibody (FITC rabbit anti-mouse IgG) (1:100 dilution) at 37°C for 1h. The cells were washed three times with PBS and counterstained with 4’, 6’-diamidino-2-phenylindole (DAPI). Coverslips were placed on slides in mounting medium and observed under a fluorescence microscope (Nikon, Japan).

### Transwell migration assay

Transwell migration assay were used to monitor the migration of THP-1 cells treated with chemerin. HUVECs were seeded in the bottom chamber of Transwell chambers and then treated with Tp0965. After incubation, THP-1 cells labeled with Calcein-AM (Sigma-Aldrich, St Louis, MO, USA) were added to upper chamber. The THP-1 cells migrated into the lower chamber were counted under a fluorescence microscope. Three replicate wells were performed per experiment. Five microscopic fields of each well were selected randomly for counting Calcein-AM labeled cells.

### Statistical analysis

Numerical data are expressed as means ± standard error. Two group comparisons were determined by Student’s *t*-test; *P* ≤ 0.05 was considered statistically significant. All experiments were performed at least in triplicate.

## Results

### Tp0965 induced expression of chemerin and ChemR23 in endothelial cells

To investigate the involvement of Tp0965 in expression of chemerin and ChemR23 in endothelial cells, we treated HUVECs with Tp0965 or vehicle (DMSO). The level of chemerin in HUVECs was elevated after stimulation using Tp0965 (Fig 1A and 1B). Likewise, expression of ChemR23 on the cell surface was significantly higher in Tp0965-treated HUVECs than in the vehicle group (Fig 1A and 1C). These results demonstrated that Tp0965 increased expression of chemerin and ChemR23, as well as secretion of chemerin, in endothelial cells.

**Fig 1.**
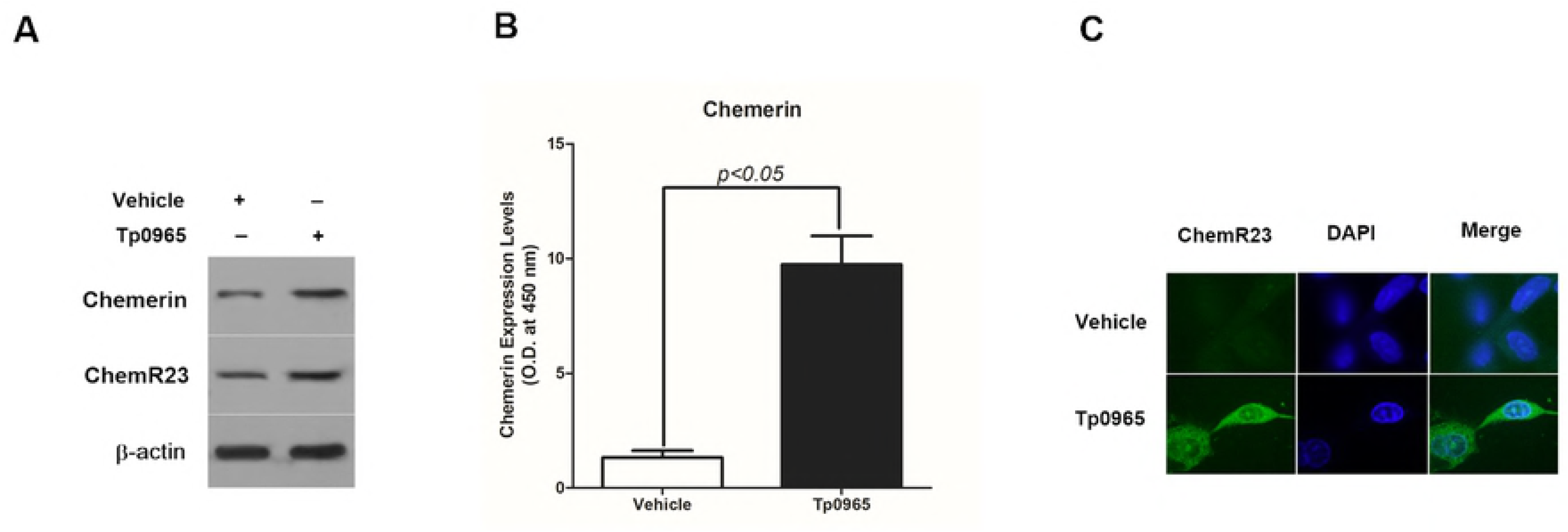
Tp0965 induced expression of chemerin and ChemR23 in endothelial cells. (A) Western blot analysis of chemerin and ChemR23 in HUVECs treated by Tp0965. (B)ELISA analysis of secreted level of chemerin in supernatant of HUVECs treated by Tp0965. Data represent mean ± SD determined from three independent experiments (n = 3), each experiment containing three technical replicates. (C) Expression of ChemR23 on HUVECs observed by confocal microscopy. HUVECs were treated by Tp0965 for 24 h. Green represents expression and distribution of ChemR23, and blue nuclear staining by DAPI.

### Toll-like receptor 2 (TLR2) is involved in the expression of chemerin

Toll-like receptor 2 (TLR2), an important member of the TLR family, recognizes bacterial lipoproteins and plays a crucial role in the host defense against bacterial infections, including those caused by *T. pallidum* [24]. Previous studies showed that TLR2 expressed on endothelial cells mediates the initiation of inflammatory signaling cascades [25]. To confirm the roles of TLR2 in the induction of chemerin expression by Tp0965, we transformed HUVECs with shRNA against TLR2 and TLR4, denoting them as HUVECs (shTLR2) and HUVECs (shTLR4), respectively. HUVECs transformed with vector alone, designated as HUVECs (V), was used as a negative control group. As expected, the expression of chemerin in HUVECs (shTLR2) was downregulated after treatment with Tp0965, but Tp0965 was unable to regulate expression of chemerin in HUVECs (shTLR4) (Fig 2), suggesting that Tp0965-induced expression of chemerin depends on TLR2 signaling.

**Fig 2.**
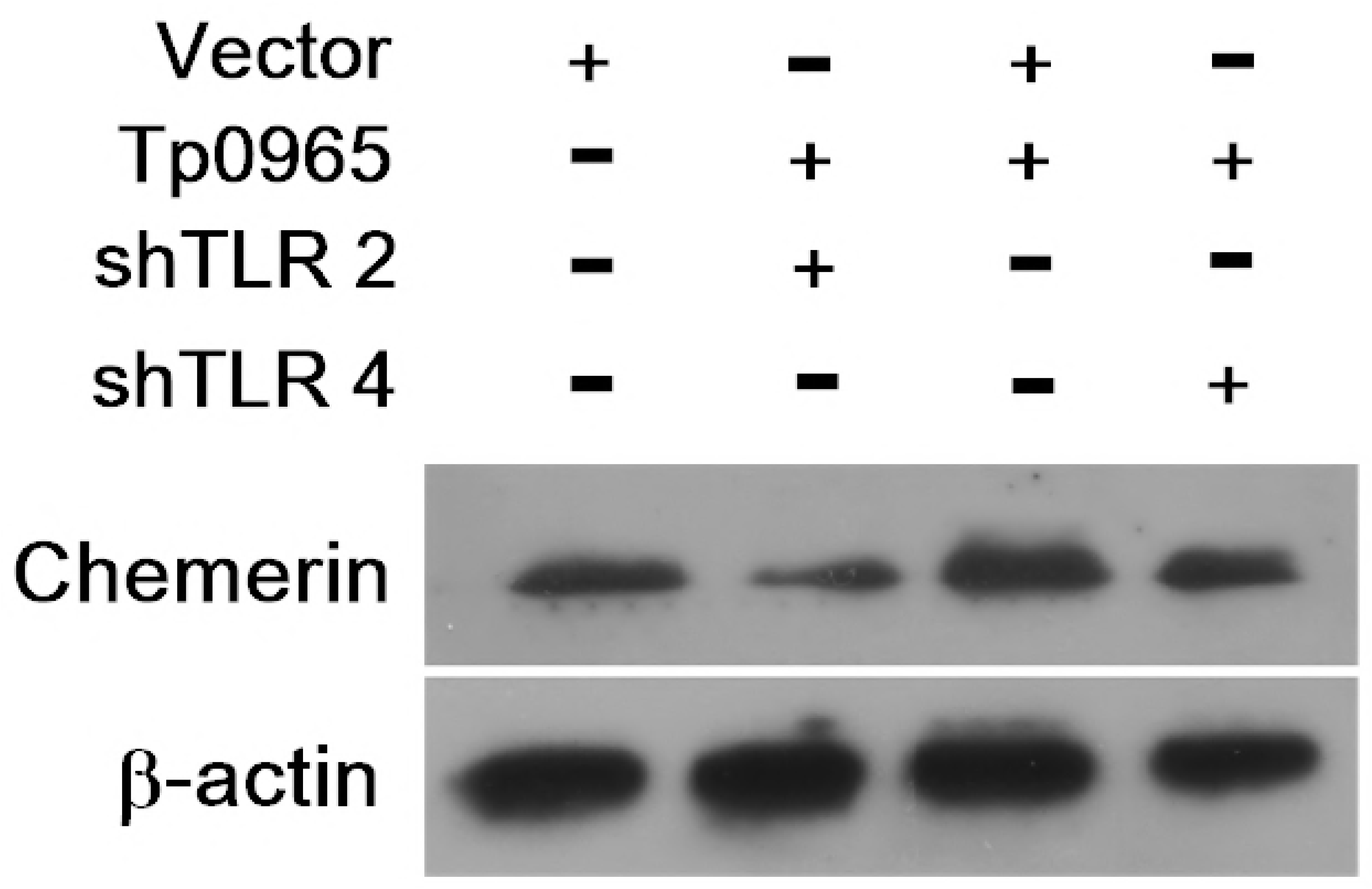
TLR2 involved in the expression of chemerin. Western blot analysis of chemerin in HUVECs. HUVECs were transformed with either shRNA of TLR2 or shRNA of TLR4, followed by treatment with Tp0965 for 24 h.

### Chemerin induces activation of endothelial cells via chemerin/ChemR23 signaling pathway

Chemerin has been showed to promote inflammation of endothelial cells. To characterize the role of chemerin in activation of endothelial cells in an autocrine/paracrine manner, we transform HUVECs with shRNA against chemerin to knockdown expression of chemerin, for use as a negative control, denoting it as HUVECs (shchemerin). Thereafter, HUVECs and HUVECs (shchemerin) were treated with Tp0965 and the supernatants were collected to stimulate HUVECs. Fig 3A showed that expression of MMP-2, ICAM-1 and E-selectin were upregulated after stimulation with the supernatant from HUVECs treated with Tp0965, similarly to the positive control with recombinant chemerin. The data demonstrated that chemerin induced by Tp0965 can promote activation of endothelial cells in an autocrine/paracrine manner.

**Fig 3.**
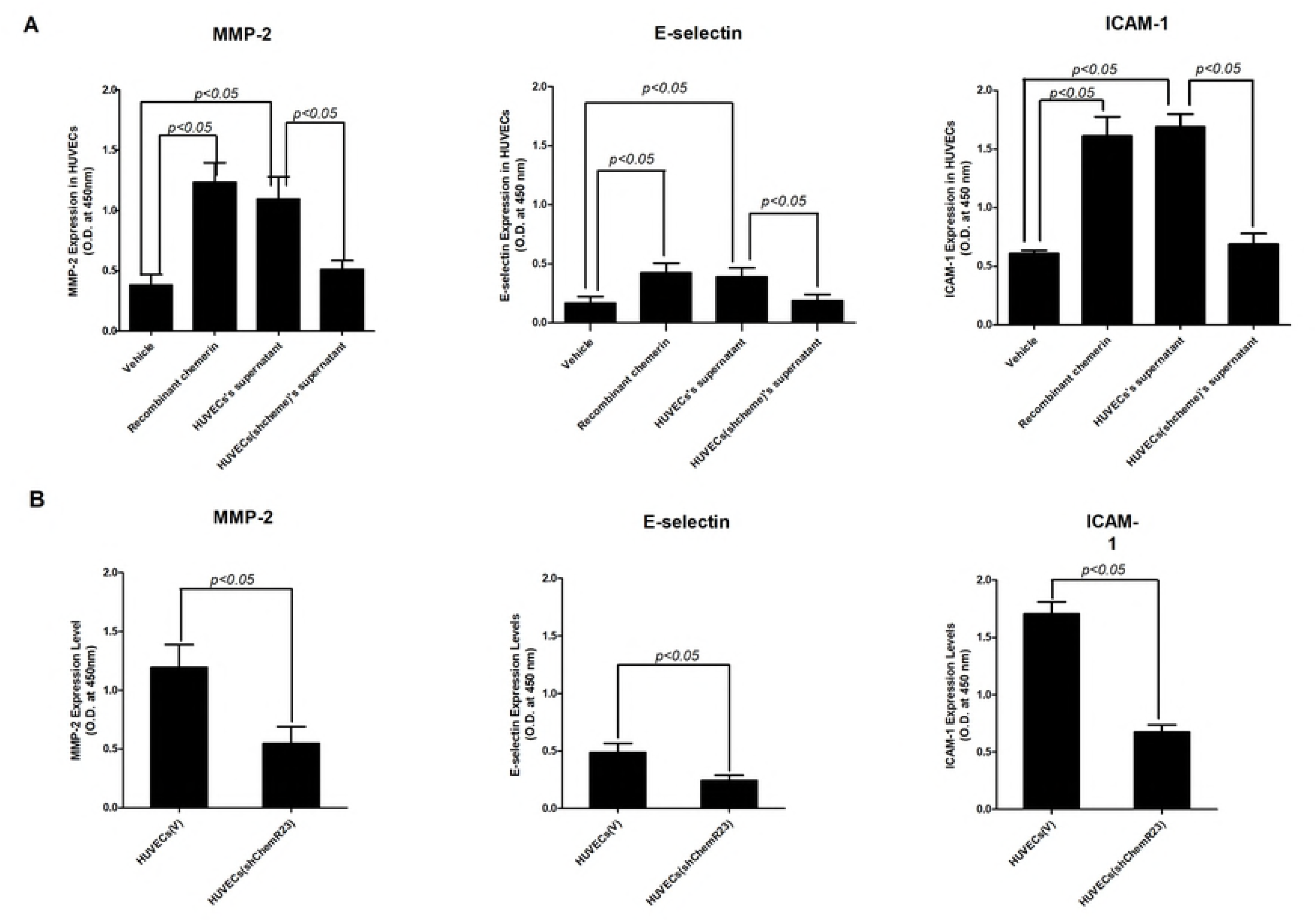
Chemerin induced activation of endothelial cells by chemerin/ChemR23 signaling pathway. (A) The secreted level of MMP-2, ICAM-1 and E-selectin in supernatant of HUVECs. HUVECs were treated with supernatant of Tp0965-treated HUVECs or recombinant chemerin for 24 h. (B) The secreted level of MMP-2, ICAM-1 and E-selectin in supernatant of HUVECs (shChemR23). HUVECs were transformed with shRNA against ChemR23, and then treated with the supernatant of Tp0965-treated HUVECs for 24 h. Data represent mean ± SD determined from three independent experiments (n = 3), each experiment containing four technical replicates. P < 0.05 for Student’s *t*-test.

Of the three chemerin receptors, only ChemR23 is responsible for the biological activities of chemerin. To confirm that chemerin promotes activation of endothelial cells through the chemerin/ChemR23 signaling pathway, we transformed HUVECs with shRNA against ChemR23, denoting it as HUVECs (shChemR23). We found expression of MMP-2, ICAM-1 and E-selectin in HUVECs(shChemR23) was decreased by supernatants from HUVECs treated with Tp0965 (Fig 3B). As expected, the role of chemerin in activation of endothelial cells depended on ChemR23.

### MAPK signaling pathway mediated chemerin induction in inflammation

MAPK signaling pathways are involved in pathophysiological processes in endothelial cells. To investigate the role of p38 MAPK signaling in activation of endothelial cells by chemerin, we measured the phosphorylation of p-JNK, p-ERK and p38 MAPK. Expression of p-ERK and p-p38 MAPK, but not p-JNK, was significantly higher in HUVECs induced with supernatants from HUVECs treated with Tp0965, as well as recombinant chemerin (Fig 4A). Next, we inhibited p-ERK and p38 MAPK with SCH772984 and SB203580, respectively, before treatment with supernatant and recombinant chemerin. Phosphorylation of p-ERK and p38 MAPK was inhibited by these compounds (Fig 4B). These results suggest that the MAPK signaling pathway is involved in regulating activation of HUVECs by chemerin.

**Fig 4.**
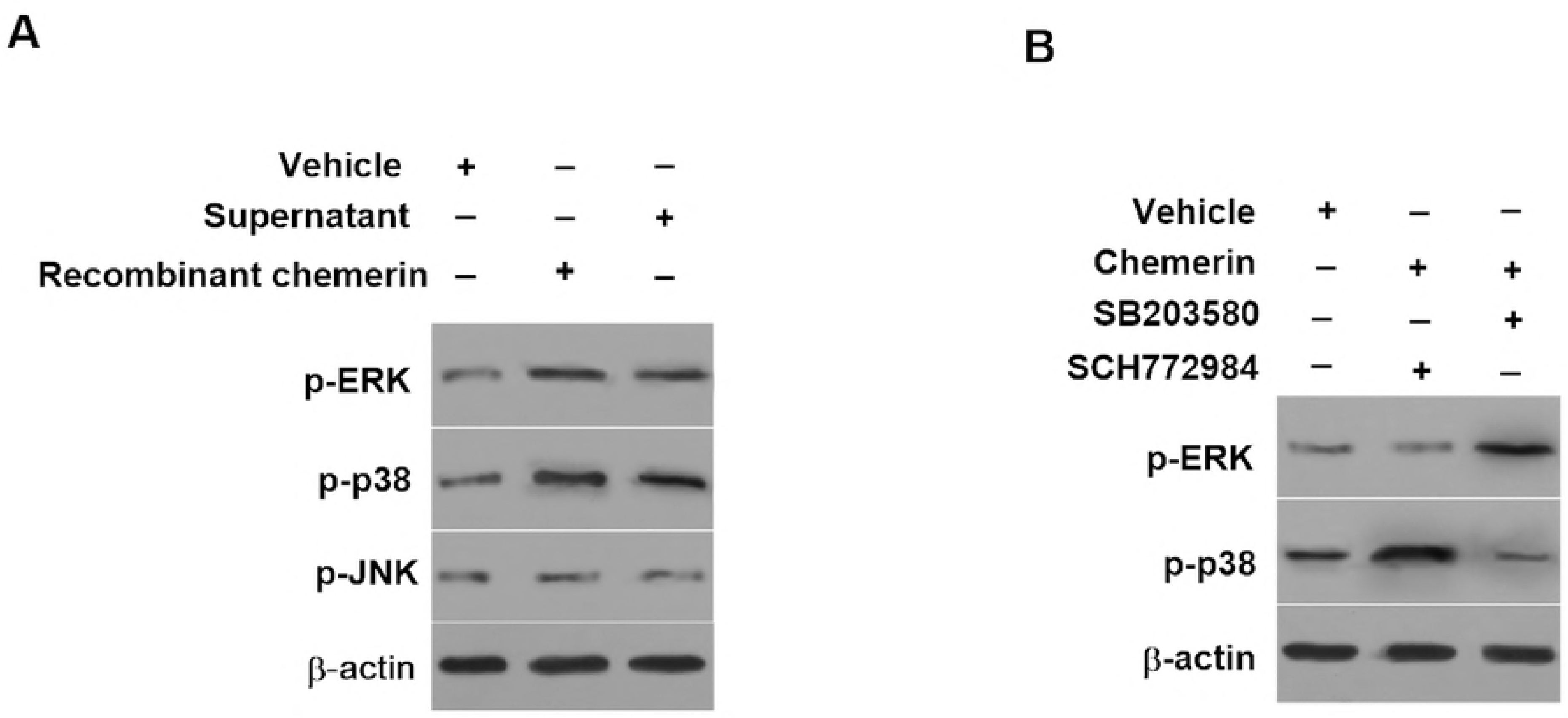
MAPK signaling pathway mediated chemerin induction in inflammation. (A) Western blot analysis of activation of p-JNK, p-ERK, and p38 MAPK in HUVECs treated with either supernatant of Tp0965-treated HUVECs or recombinant chemerin for 24 h. (B) Western blotting analysis of activation of p-ERK and p38 MAPK in HUVECs. HUVECs were pre-incubated with SCH772984 and SB203580, respectively, and then treated with above supernatant and recombinant chemerin for 24 h.

### Tp0965 promotes migration of monocytes through the chemerin/ChemR23 signaling pathway

Chemerin/ChemR23 regulate inflammation, including leukocyte recruitment and migration through the extracellular matrix to the site of inflammation [26]. To characterize the effect of Tp0965-induced chemerin on monocytes recruitment and migration to endothelial cells, we pre-incubated HUVECs seeded in the bottom chambers of Transwell with supernatants from HUVECs treated with Tp0965 or recombinant chemerin. We then added Calcein-AM-labeled THP-1 cells to Transwell insert and measured the migration of THP-1 cells in response to chemerin. In a previous study, we showed Tp0965 significantly upregulated expression of soluble MCP-1, known as chemotactic factor to monocytes [13]. To inhibit the effect of MCP-1, every bottom chamber contained monoclonal antibodies against MCP-1 before the addition of THP-1 cells. Fig 5 showed that both supernatant and recombinant chemerin had the ability to sustain the migration of THP-1 cells. Next, we knocked down ChemR23 expression with shRNA and added these knockdown cells to the Transwell insert. THP-1 cells (shChemR23) were unable to migrate in response to either the supernatant or recombinant chemerin. Conversely, no effect was observed with vehicle alone (Fig 5). These results formally proved that Tp0965-induced chemerin promoted monocytes migration via chemerin/ChemR23 signaling pathway.

**Fig 5.**
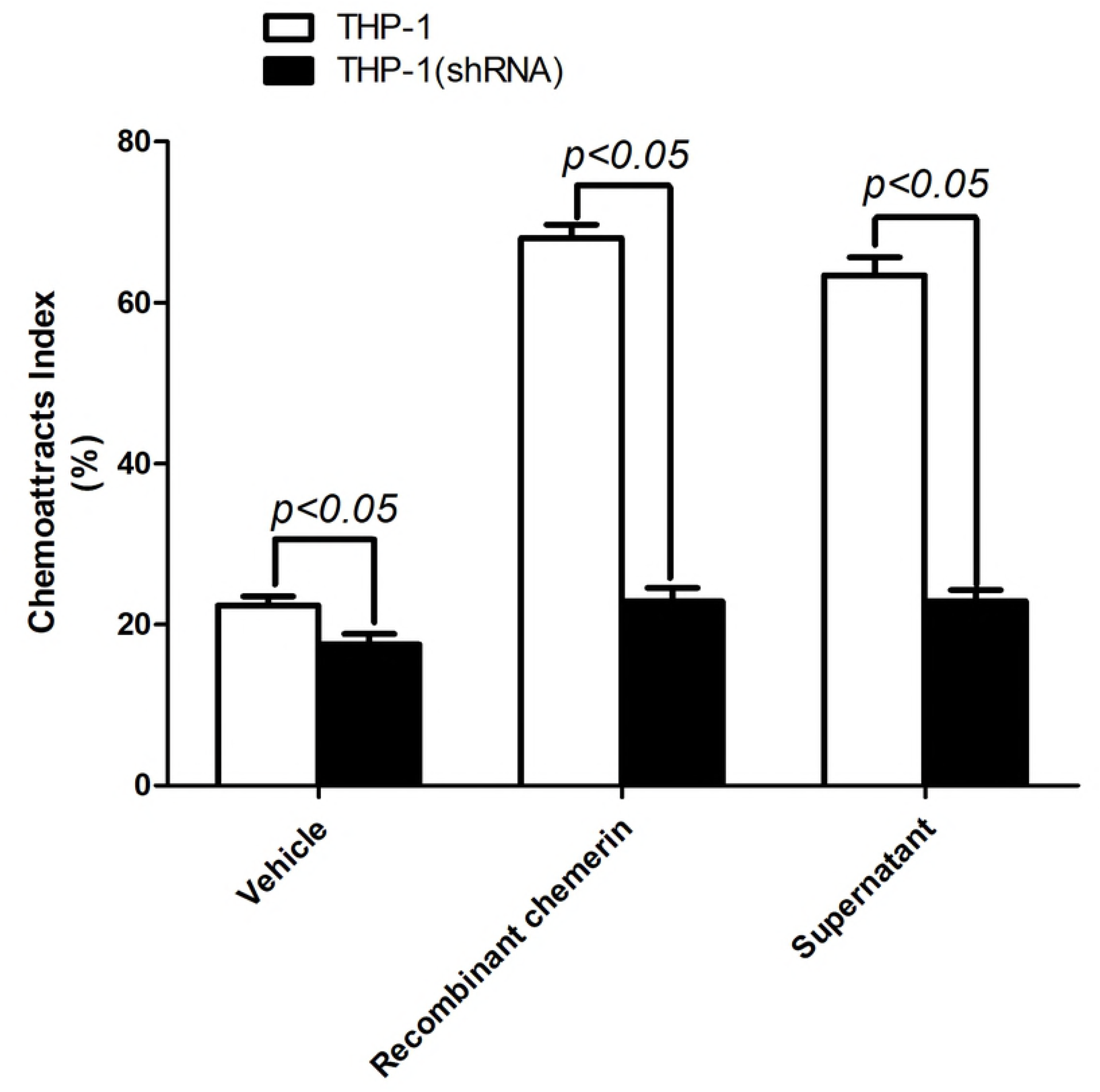
Tp0965 promoted migration of monocytes through chemerin/ChemR23 signaling pathway. Transwell migration assays were performed in HUVECs. HUVECs were seeded in Transwell chambers, and then treated with either supernatants or recombinant chemerin. THP-1 cells migrated into the lower chamber was counted. On the other experiment, THP-1 cells were transformed with shChemR23 before being added in upper chambers, and then were counted. P < 0.05 for Student’s *t-*test.

### RhoA/ROCK signaling mediates migration of monocytes

Macrophages migration to sites of inflammation is dependent on the RhoA/ROCK signaling pathway [27]. This signaling pathway is also required for chemerin-mediated chemotaxis of lymphocytes [28]. Previously, we reported that RhoA in HUVECs is activated following treatment with Tp0965 [13]. To further explore this issue, we investigated whether RhoA was activated in HUVECs stimulated with supernatants of Tp0965-treated HUVECs or recombinant chemerin. Indeed, RhoA was activated following either treatment (Fig 6A). We then examined the effect of ROCK on monocyte migration. For this purpose, we pretreated HUVECs with Y-27632, an inhibitor of ROCK. In addition, cells were treated with monoclonal antibodies against MCP-1 to eliminate the effect of this cytokine on monocyte migration. Chemotaxis assays revealed no significant changes relative to vehicle, indicating that ROCK activity required for monocyte migration (Fig 6B). These data suggest that the RhoA/ROCK signaling pathway is involved in Tp0965-induced monocytes migration in response to chemerin/ChemR23.

**Fig 6.**
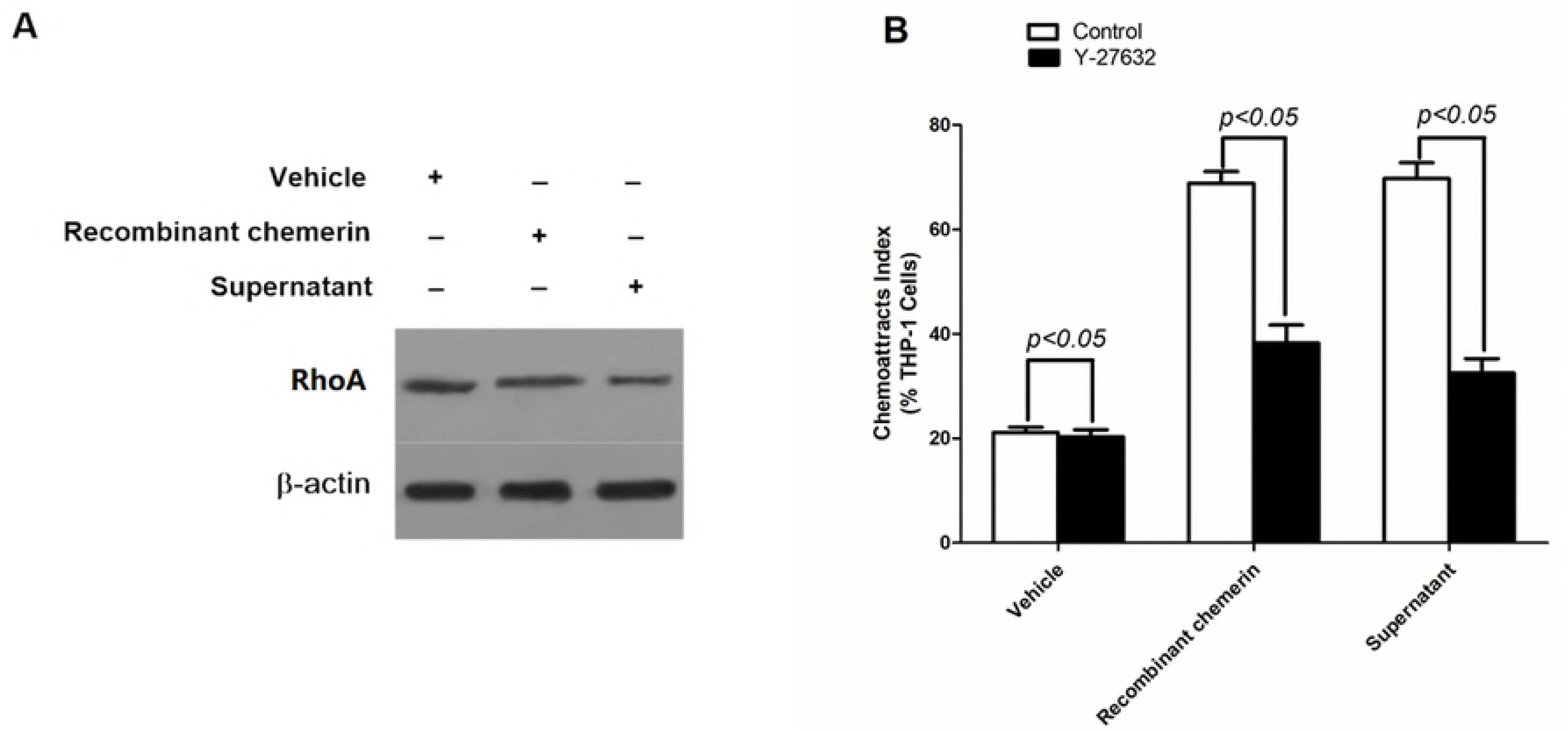
RhoA/ROCK signaling mediated migration of monocytes. (A) Western blot analysis of activation of RhoA in HUVECs treated with either supernatant of Tp0965-treated HUVECs or recombinant chemerin for 24 h. (B) Transwell migration assays were performed in HUVECs. HUVECs were pretreated with Y-27632, and then treated with either supernatants or recombinant chemerin. THP-1 cells migrated into the lower chamber was counted. P < 0.05 for Student’s *t*-test.

## Discussion

Inflammation response was emphasized to be responsible for syphilis pathogenesis during the invasion and persistence of *T. pallidum.* According to a recent proposed schema [29], treponemal lipoproteins created an proinflammatory backdrop and then an adaptive immune response stepped-up as treponemal antigens are processed and presented to naive T cells. Consequently, vasculitis was invoked by recruitment of immune cells, and led to *T. pallidum* dissemination through disrupting endothelial barrier. Thus, the investigation focus on endothelium damage will contribute to declare the immunopathogenesis of syphilis.

Chemerin is well characterized by acting as a chemoattractant for immune cells expressing ChemR23. Thereby, chemerin plays a pro-inflammatory role via recruiting these immune cells to sites of inflammation and tissue damage [30-31]. In a recent study, we observed a significant increase in serum level of chemerin in HUVECs following treatment with *T. pallidum* Nichols strain (data not shown). On the basis of this finding, we hypothesized that Tp0965 induces the dysfunction of endothelial cells through chemerin signaling. To test the hypothesis, in this present study, we investigated the effect of Tp0965 on chemerin expression in vascular endothelial cells. The results revealed that Tp0965 can significantly increase the expression of chemerin in HUVECs. TLR4 and very low levels of TLR2 are expressed in normal endothelial cells [32]. However, TLR2 expression is dramatically increased upon stimulation with inflammatory factors. In contrast to TLR4, which potentiates macrophage responses during inflammation [33], TLR2 is expressed more broadly in vascular cells and promotes endothelial dysfunction [34]. To further characterize the relationship between TLR2 and chemerin, we knocked down TLR2 and TLR4 of HUVECs with shRNA. Expression of chemerin was downregulated after Tp0965 treatment in HUVECs (shTLR2), but not in HUVECs (shTLR4). These results suggested that TLR2 signaling is involved in Tp0965-induced chemerin expression. Several studies demonstrated that TLR2 promotes endothelial dysfunction by initiating inflammatory signaling cascades [35-36]. However, further study is necessary to determine how TLR2 is involved in expression of chemerin in endothelial cells.

As a mediator of vascular inflammation, chemerin is involved in angiogenesis, atherosclerosis, and adipocyte metabolism, ultimately leading to the development of cardiovascular diseases [37]. In addition, chemerin may induce activation of vascular endothelial cells, characterized by expression of cytokines including adhesion molecules and endothelial gelatinases, thereby promoting the growth and remodeling of blood vessels [38-40]. Accordingly, we investigated the effects of chemerin on endothelial cells. The results revealed that chemerin significantly upregulated expression of ICAM-1, E-selectin, and MMP-2 in HUVECs. ICAM-1 and E-selectin are markers of endothelial cells activation, and can induce rapid adhesion of leukocytes to vascular endothelium and recruit these cell to sites of inflammation. On the other hand, MMP-2 contributes to dysfunction of the endothelial barrier. Together, these data suggest a pro-inflammatory role for chemerin in endothelial dysfunction. However, several studies suggested that chemerin plays anti-inflammatory roles in vascular endothelial cells [41-42]. Yamawaki *et al.* reported that chemerin prevents TNF-α-induced VCAM-1 expression and monocytes adhesion in vascular endothelial cells [42]. This discrepancy may be due to difference in how chemerin protein is processed. Chemerin is initially secreted as pro-chemerin, and is activated by cleavage by proteases such as inflammatory serine proteases tryptase, elastase, and plasmin [43]. The variant cleavages give rise to different chemerin functions, potentially explaining the inconsistent results reported by previous studies.

In this study, we observed upregulation of ChemR23 expression in HUVECs after treatment with Tp0965. ChemR23 is an orphan G protein-coupled receptor also known as a chemokine-like receptor 1 (CMKLR1); the other two receptors of chemerin are chemokine C-C motif receptor-like 2 (CCRL-2) and G protein-coupled receptor 1 (GPR 1). Among the three receptors, ChemR23 is predominantly responsible for the function of chemerin in inflammation [44]. Consistent with previous research, we found that both endothelial-derived and recombinant chemerin increased the expression of MMP-2, ICAM-1 and E-selectin in HUVECs. However, knockdown of the ChemR23 gene in HUVECs partly inhibited the pro-inflammatory function of chemerin. These findings suggest that chemerin affects endothelial cells through the chemerin/ChemR23 signaling way in an autocrine/paracrine manner.

The MAPK signaling pathways have three major components: ERK, p38, and JNK. These proteins exist in most immune cells and are involved in both physiological and pathophysiological processes. Previous research demonstrated that chemerin effects on monocyte-endothelial adhesion via the MAPK and PI3K/Akt pathways [45]. In the present study, we observed that chemerin induced the phosphorylation of ERK MAPK, and p38 MAPK in HUVECs. This ability of chemerin was abolished by treatment with p38 MAPK inhibitor. However, chemerin had no effect of the phosphorylation of p-JNK. These results agreed with findings reported in recent studies [46]. Kaur *et al*. demonstrated that chemerin dose-dependently activates ERK MAPK and p38 MAPK in endothelial cells [40]. Our data suggest that both recombinant and Tp0965-induced chemerin promote the activation of HUVECs through the MAPK signaling pathway.

Chemerin was originally described as a chemoattractant that promotes recruitment of ChemR23-expressing immune cells, including immature dendritic cells (DCs), natural killer (NK) cells, and macrophages, to sites of inflammation [47-49]. In addition, a recent study showed that endothelial cells expressing chemerin can promote DCs transmigration across endothelial cell barriers, mediated by ChemR23/chemerin signaling [22]. Conversely, DCs can induce endothelial cells to express chemerin protein. As reported previously, in this study, we observed migration of THP-1 cells to HUVECs following treatment with Tp0965-induced chemerin. Furthermore, the migration of THP-1 cells was inhibited by knockdown of ChemR23. These results suggest that Tp0965-induced chemerin promotes monocytes migration through the chemerin/ChemR23 signaling pathway. Migration and retention of leukocytes during inflammation induced by chemerin are thought to contribute to the onset and development of inflammatory diseases, as reported in atherosclerosis and arthritis [50-51]. During the development of atherosclerosis, macrophages from the surrounding intima and adventitia migrate into the atherosclerotic lesion and adhere to extracellular matrix proteins. The adhesion of macrophages might decrease plaque stability by upregulating MMPs expression [52-53]. In this study, we also observed upregulation of MMP-2 expression in endothelial cells. However, the role of MMP-2 in endothelial cell dysfunction induced by *T. pallidum* warrants further exploration.

In a previous study, we found that the RhoA/ROCK signaling pathway is involved in increasing endothelial permeability and promotes transendothelial migration of monocytes [13]. Other groups also reported that the RhoA/ROCK signaling pathway plays important roles in the pathogenesis of cardiovascular disease [54]. In addition, Rourke *et al*. showed that RhoA knockdown, or pharmacological inhibition of RhoA or ROCK, blocks chemerin-induced activation of serum response factor [28]. In this study, we observed activation of RhoA in HUVECs after stimulation with Tp0965-induced chemerin. Furthermore, when we inhibited ROCK with Y-27632, we found that monocyte migration to HUVECs was blocked. These findings verified that Tp0965-induced monocytes migration in response to chemerin/ChemR23 depends on the activity of the RhoA/ROCK signaling pathway.

This study has one major limitation. We showed that chemerin induced by Tp0965 affects the function of endothelial cells via the chemerin/ChemR23 signaling pathway *in vitro*. However, it remains unknown whether chemerin can induce dysfunction of vascular endothelium *in vivo*. Unfortunately, no animal model system that can mimic the behavior of *T. pallidum* in the host body is currently available. Notably in this regard, a recent study reported that the zebrafish model was successfully used to observe the angiogenic activity of TpF1 of *T. pallidum* [55]. Accordingly, we anticipate that the zebrafish model could be used to study the angiogenic effect of Tp0965 on endothelial cells *in vivo*.

In summary, we showed that Tp0965-induced chemerin promotes dysfunction of endothelial cells, dependent on the activation of MAPK and RhoA/ROCK signaling pathways. These results provide further insight into the effect of Tp0965 on the function of endothelial cells, suggesting the role of predicted membrane proteins of *T. pallidum* in the immunopathogenesis of vascular inflammation of syphilis.

## Acknowledgements

The authors thank Dr. Wenlong Hu and Ms. Bufang Xu for technical assistance.

## References

1. World Health Organization. Global incidence and prevalence of selected sexually transmitted infections– 2008. Geneva, 2012. Available at: http://sci-hub.tw/ http://www.who.int/reproductivehealth/publications/rtis/stisestimates/en/

2. Byard RW. Syphilis-Cardiovascular Manifestations of the Great Imitator. J Forensic Sci 2017; doi:10.1111/1556-4029.13709.

3. Roberts WC, Bose R, Ko JM, Henry AC, Hamman BL. Identifying cardiovascular syphilis at operation. Am J Cardiol 2009; 104:1588–1594.

4. Deka RK, Brautigam CA, Liu WZ, Tomchick DR, Norgard MV. Evidence for Posttranslational Protein Flavinylation in the Syphilis Spirochete Treponema pallidum: Structural and Biochemical Insights from the Catalytic Core of a Periplasmic Flavin-Trafficking Protein. MBio 2015; 6: e00519–15.

5. Liu J, Howell JK, Bradley SD, Zheng Y, Zhou ZH, Norris SJ. Cellular architecture of Treponema pallidum: novel flagellum, periplasmic cone, and cell envelope as revealed by cryo electron tomography. J Mol Biol 2010; 403:546–561.

6. McGill MA, Edmondson DG, Carroll JA, Cook RG, Orkiszewski RS, Norris SJ. Characterization and serologic analysis of the Treponema pallidum proteome. Infect Immun 2010; 78:2631–2643.

7. Osbak KK, Houston S, Lithgow KV, Meehan CJ, Strouhal M, Šmajs D, et al. Characterizing the Syphilis-Causing Treponema pallidum ssp. pallidum Proteome Using Complementary Mass Spectrometry. Van Ostade X. Kenyon CR. Van Raemdonck GA. PLoS Negl Trop Dis 2016; 10(9): e0004988.

8. Luo X, Zhang X, Zhao T, Zeng T, Liu W, Deng M, Zhao F. A preliminary study on the proinflammatory mechanisms of Treponema pallidum outer membrane protein Tp92 in human macrophages and HMEC-1 cells. Microb Pathog 2017; 110:176–183.

9. Houston S, Russell S, Hof R, et al. The multifunctional role of the pallilysin-associated Treponema pallidum protein, Tp0750, in promoting fibrinolysis and extracellular matrix component degradation. Mol Microbiol. 2014; 91: 618–634.

10. Runina AV, Starovoitova AS, Deryabin DG, Kubanov AA. Evaluation of the Recombinant Protein Tp0965 of Treponema Pallidum as Perspective Antigen for the Improved Serological Diagnosis of Syphilis. Vestn Ross Akad Med Nauk 2016; 2:109–113.

11. Long FQ, Zhang JP, Shang GD, Shang SX, Gong KL, Wang QQ. Seroreactivity and immunogenicity of Tp0965, a hypothetical membrane protein of Treponema pallidum. Chin Med J (Engl) 2012; 125(11):1920–1924.

12. Bonetti PO, Lerman LO, Lerman A. Endothelial dysfunction: a marker of atherosclerotic risk. Arterioscler Thromb Vasc Biol 2003; 23: 168–175.

13. Riley BS, Oppenheimer-Marks N, Hansen EJ, Radolf JD, Norgard MV. Virulent Treponema pallidum activates human vascular endothelial cells. J Infect Dis 1992; 165(3):484–493.

14. Kao WA, Pětrošová H, Ebady R, Lithgow KV, Rojas P, Zhang Y, et al. Identification of Tp0751 (Pallilysin) as a Treponema pallidum Vascular Adhesin by Heterologous Expression in the Lyme disease Spirochete. Sci Rep 2017; 7(1):1538.

15. Zhang RL, Zhang JP, Wang QQ. Recombinant treponema pallidum protein Tp0965 activates endothelial cells and increases the permeability of endothelial cell monolayer. PLoS ONE 2014; 12: e115134.

16. Zhang RL, Zhang JP, Wang QQ, Yang LJ. Tp17 member protein of treponema pallidum activates endothelial cell in vitro. Int Immunopharmacol 2015; 25:538– 544.

17. Lin S, Teng J, Li J, Sun F, Yuan D, Chang J. Association of Chemerin and Vascular Endothelial Growth Factor (VEGF) with Diabetic Nephropathy. Med Sci Monit 201;2: 3209–3214.

18. Neves KB, Nguyen Dinh Cat A, Lopes RA, et al. Chemerin Regulates Crosstalk Between Adipocytes and Vascular Cells Through Nox. Hypertension 2015; 66: 657–666.

19. Monnier J, Lewén S, O’Hara E, Huang K, Tu H, Butcher EC, Zabel BA. Expression, regulation, and function of atypical chemerin receptor CCRL2 on endothelial cells. J Immunol 2012; 189:956–967.

20. Salama FE, Anass QA, Abdelrahman AA, Saeed EB. Chemerin: A biomarker for cardiovascular disease in diabetic chronic kidney disease patients. Saudi J Kidney Dis Transpl 2016; 27: 977–984.

21. Akamata K, Asano Y, Taniguchi T, et al. Increased expression of chemerin in endothelial cells due to Fli1 deficiency may contribute to the development of digital ulcers in systemic sclerosis. Rheumatology (Oxford) 2015; 54:1308–1316.

22. De Palma G, Castellano G, Prete AD, et al. The possible role of ChemR23/Chemerin axis in the recruitment of dendritic cells in lupus nephritis. Kidney Int 2011; 79:1228–1235.

23. Albanesi C, Scarponi C, Pallotta S, et al. Chemerin expression marks early psoriatic skin lesions and correlates with plasmacytoid dendritic cell recruitment. J Exp Med 2009; 206:249–258.

24. Xie Y, Xu M, Xiao Y, et al. Treponema pallidum flagellin FlaA2 induces IL-6 secretion in THP-1 cells via the Toll-like receptor 2 signaling pathway. Mol Immunol 2016; 81:42–51.

25. Benhamou Y, Bellien J, Armengol G, et al. Role of Toll-like receptors 2 and 4 in mediating endothelial dysfunction and arterial remodeling in primary arterial antiphospholipid syndrome. Arthritis Rheumatol 2014; 66:3210–3220.

26. Bondue B, Wittamer V, Parmentier M. Chemerin and its receptors in leukocyte trafficking, inflammation and metabolism. Cytokine Growth Factor Rev 2011; 22:331–338.

27. Van Goethem E, Poincloux R, Gauffre F, Maridonneau-Parini I, Le Cabec V. Matrix architecture dictates three-dimensional migration modes of human macrophages: differential involvement of proteases and podosome-like structures. J Immunol 2010; 184:1049–1061.

28. Rourke JL, Dranse HJ, Sinal CJ. CMKLR1 and GPR1 mediate chemerin signaling Rourke through the RhoA/ROCK pathway. Mol Cell Endocrinol 2015; 417:36–51.

29. Salazar JC, Hazlett KR, Radolf JD. The immune response to infection with Treponema pallidum, the stealth pathogen. Microbes Infect 2002; 4(11):1133–1140.

30. Landgraf K, Friebe D, Ullrich T, et al. Chemerin as a mediator between obesity and vascular inflammation in children. J Clin Endocrinol Metab 2012; 97: E556-564.

31. Skrzeczyńska-Moncznik J, Wawro K, Stefańska A, et al. Potential role of chemerin in recruitment of plasmacytoid dendritic cells to diseased skin. Biochem Biophys Res Commun 2009; 380:323–327.

32. Dunzendorfer S, Lee HK, Tobias PS. Flow-dependent regulation of endothelial Toll-like receptor 2 expression through inhibition of SP1 activity. Circ Res 2004; 95: 684–691.

33. Hachiya R, Shiihashi T, Shirakawa I, et al. The H3K9 methyltransferase Setdb1 regulates TLR4-mediated inflammatory responses in macrophages. Sci Rep 2016; 6:28845.

34. Diesel B, Ripoche N, Risch RT, Tierling S, Walter J, Kiemer AK. Inflammation-induced up-regulation of TLR2 expression in human endothelial cells is independent of differential methylation in the TLR2 promoter CpG island. Innate Immun 2012; 18:112–123.

35. Xu Y, Zhou Y, Lin H, Hu H, Wang Y, Xu G. Toll-like receptor 2 in promoting angiogenesis after acute ischemic injury. Int J Mol Med 2013; 31:555–560.

36. Wilhelmsen K, Mesa KR, Lucero J, Xu F, Hellman J. ERK5 protein promotes, whereas MEK1 protein differentially regulates, the Toll-like receptor 2 protein-dependent activation of human endothelial cells and monocytes. J Biol Chem 2012; 287:26478–26494.

37. Bonomini M, Pandolfi A. Chemerin in renal dysfunction and cardiovascular disease. Vascul Pharmacol 2016; 77:28–34.

38. Ferland DJ, Watts SW. Chemerin: A comprehensive review elucidating the need for cardiovascular research. Pharmacol Res 2015; 99:351–361.

39. Bozaoglu K, Curran JE, Stocker CJ, et al. Chemerin, a novel adipokine in the regulation of angiogenesis. J Clin Endocrinol Metab 2010; 95: 2476–2485.

40. Kaur J, Adya R, Tan BK, Chen J, Randeva HS. Identification of chemerin receptor (ChemR23) in human endothelial cells: chemerin-induced endothelial angiogenesis. Biochem Biophys Res Commun 2010; 391:1762–1768.

41. Wang L, Yang T, Ding Y, Zhong Y, Yu L, Peng M. Chemerin plays a protective role by regulating human umbilical vein endothelial cell-induced nitric oxide signaling in preeclampsia. Endocrine 2015; 48:299–308.

42. Yamawaki H, Kameshima S, Usui T, Okada M, Hara Y. A novel adipocytokine, chemerin exerts anti-inflammatory roles in human vascular endothelial cells. Biochem Biophys Res Commun 2012; 423:152–157.

43. Du XY, Leung LL. Proteolytic regulatory mechanism of chemerin bioactivity. Acta Biochim Biophys Sin (Shanghai) 2009; 41:973–979.

44. Mattern A, Zellmann T, Beck-Sickinger AG. Processing, signaling, and physiological function of chemerin. IUBMB Life 2014; 66:19–26.

45. Dimitriadis GK, Kaur J, Adya R, et al. Chemerin induces endothelial cell inflammation: activation of nuclear factor-kappa beta and monocyte-endothelial adhesion. Oncotarget 2018; 9:16678–16690.

46. Chua SK, Shyu KG, Lin YF, et al. Tumor Necrosis Factor-Alpha and the ERK Pathway Drive Chemerin Expression in Response to Hypoxia in Cultured Human Coronary Artery Endothelial Cells. PLoS ONE 2016; 11: e0165613.

47. Yin Q, Xu X, Lin Y, Lv J, Zhao L, He R. Ultraviolet B irradiation induces skin accumulation of plasmacytoid dendritic cells: a possible role for chemerin. Autoimmunity 2014; 47:185–192.

48. Parolini S, Santoro A, Marcenaro E, et al. The role of chemerin in the colocalization of NK and dendritic cell subsets into inflamed tissues. Blood 2007; 109:3625–3632.

49. Maheshwari A, Kurundkar AR, Shaik SS, et al. Epithelial cells in fetal intestine produce chemerin to recruit macrophages. Am J Physiol Gastrointest Liver Physiol 2009; 297: G1–G10.

50. McNeill E, Iqbal AJ, Patel J, et al. Contrasting in vitro vs. in vivo effects of a cell membrane-specific CC-chemokine binding protein on macrophage chemotaxis. J Mol Med (Berl) 2014; 92: 1169–1178.

51. Mariani F, Roncucci L. Chemerin/ChemR23 axis in inflammation onset and resolution. Inflamm Res 2015; 64: 85–95.

52. Johnson JL. Matrix metalloproteinases: influence on smooth muscle cells and atherosclerotic plaque stability. Expert Rev Cardiovasc Ther 2007; 5:265–282.

53. Newby AC. Metalloproteinase expression in monocytes and macrophages and its relationship to atherosclerotic plaque instability. Arterioscler Thromb Vasc Biol 2008; 28: 2108–2114.

54. Zhang H, Shi L, Ren GQ, et al. Dihydrotestosterone modulates endothelial progenitor cell function via RhoA/ROCK pathway. Am J Transl Res 2016; 8:4300– 4309.

55. Pozzobon T, Facchinello N, Bossi F, et al. Treponema pallidum (syphilis) antigen TpF1 induces angiogenesis through the activation of the IL-8 pathway. Sci Rep 2016; 6:18785.

